# Light and substrate composition control root exudation rates at the initial stages of soilless lettuce cultivation

**DOI:** 10.1101/2024.05.29.596433

**Authors:** Brechtje R. de Haas, Eva Oburger, Marie-Christine Van Labeke, Emmy Dhooghe, Danny Geelen

**Author notes:** Corresponding author: D. Geelen, 0032 (0)9 264 60 76.

## Abstract

Plant root exudation is an inherent metabolic process that enhances various functions of the root system like the mobilization of nutrients and interactions with surrounding microbial communities. The role and extent of root exudation in soilless crop production is poorly investigated. Here, we analyzed soilless lettuce and show that the root exudation rate declines with plant age. Furthermore, the impact of light quality and substrate was assessed by growing soilless lettuce in 100% red light (660 nm), 100% blue light (450 nm), and white light (full-light spectrum) and in 100% perlite, 100% potting soil, or mixtures of both materials. Root exudates were collected at 10, 17 and 24 days after transplanting. The total carbon root exudation rate was influenced by light conditions and substrate composition at the earliest timepoint of the culture but not at later growth stages. The total carbohydrate exudation rate was significantly higher under pure blue and red light compared to white light. The impact of light depended on the presence of perlite in the substrate. The total phenolic compound exudation rate was most strongly influenced by the substrate composition and reached the highest level in either pure potting soil or pure perlite. These findings underscore the importance of root exudation during the initial stages of development. Light and growing media influence the exudation rate at this early stage, suggesting that exudation is an adaptive process of the soilless lettuce culture.

**HIGHLIGHT:** At an initial stage of development, soilless lettuce exudation rates are high and modulated by light and substrate composition, whereas older plants show lower rates that are not influenced by these environmental conditions.

## INTRODUCTION

Root exudation entails the chemical communication between the plant root system and its environment. The molecules involved are defined as “plant-derived primary and secondary metabolites of both low (< 1000 Da; e.g. sugars, organic acids, phenolics, vitamins) and high molecular weight (> 1000 Da; e.g. enzymes, mucilage)” (Oburger and Jones, 2018). Root exudation is not a static process, but is dynamic. Both the exudation rate and the exudation profile depend on multiple factors, a few of which are described below.

A factor that influences root exudation is the physiological age of the plant. For *Arabidopsis thaliana*, for example, the exudation of sugars and sugar alcohols decreases, while the exudation of phenolics and amino acids increases throughout their lifecycle (Chaparro *et al*., 2013). *Zea mays* shows a decline in its exudation rate of carbohydrates, amino acids, proteins and phenolics starting from the four-leaf stage up to early ripening (Santangeli *et al*., 2024). Phytosiderophore exudation of *Triticum aestivum* reduced also over time (Oburger *et al*., 2014). However, phenolic exudation of *Sorghum halepense* varied over time depending on the specific compound that was measured (Huang *et al*., 2015).

The environment in which roots grow has a profound impact on the rate and composition of exudate produced. For instance, soil type was shown to affect the exudation pattern of *Lactuca sativa*, secreting more sugars, amino acids, and organic acids, when grown in diluvial sand compared to alluvial loam (Neumann *et al*., 2014). *Zea mays* root exudation increases when sand is mixed into the soil, suggesting that soil structural properties are important (Fischer *et al*., 2010). Soil, however, has also physico-chemical properties that can affect root exudation. These properties include water holding capacity, soil pore volume, particle size and particle surface charge, nutrient availability, organic carbon content, and pH (Liang *et al*., 2013; Sardans *et al*., 2023). Soil pore volume can increase rhizodeposition of ^14^C, depending on species (Lucas *et al*., 2023). Particle size can impact root exudation by changing the root architecture. A higher particle size is related to more mechanical impedance. This may result in more lateral roots, thus increasing root exudation (Groleau-Renaud *et al*., 1998). Particle surface charge affects the adsorption to the soil, so the exudation profile of plants grown in clay differs from those grown in sand or glass (Sasse *et al*., 2020).

Because of its link with photosynthesis, light intensity has been viewed as an important environmental factor influencing root exudation. A study applying different light intensities has been shown to modulate the concentration of different amino acids in exudates from tomato and clover (Rovira, 1959). In seagrass, low light intensities reduced root exudation of total dissolved N and altered the composition of the root microbiome (Martin *et al*., 2018). Root exudation has also been shown to follow a diurnal pattern in some species and this was found to be associated with the acquisition of Fe through the secretion of phytosiderophores (Bernards *et al*., 2014; Selby-Pham *et al*., 2017). Increasing light intensity from 150 to 200 µmol m^-2^ s^-1^ reduced total C exudation in *Lactuca sativa* (Zhou *et al*., 2022). Whether this is a direct effect is unclear as high light intensity stimulates plant development and physiologically ages the plant more quickly.

The impact of different light wavelengths on root exudation has received very little attention. The lateral irradiation with far-IR light (wavelength > 10.000 nm) of seedlings of *Avena fatua* has been shown to increase the secretion of sesquiterpenes (Pomilio *et al*., 2000). The root exudation profile of *Solanum lycopersicum* changed with red/far-red ratio (Guo *et al*., 2024). Pure red and blue LED light combined stimulated a higher exudation compared to plants grown in broad-spectrum light (Zhou *et al*., 2022). UV-B light was shown to have contrasting effects on root exudation composition in different plant species (Rinnan *et al*., 2006; He *et al*., 2016).

Given these studies, we hypothesized that light intensity and quality are important factors influencing root exudation and possibly also the root microbiome. As indoor farming is ameanable to light quality control, the relationship between light and root exudation may provide an approach to control root exudation. Therefore, we analyzed the impact of light on lettuce grown in soilless conditions. We found that the effect of light quality and the type of substrate was most pronounced during the initial culturing stage. Our study further reveals that substrate composition is a key factor that determines how light quality influences root exudation.

## MATERIALS AND METHODS

### Plant material and growth conditions

*Lactuca sativa L*. (lettuce), cultivar ‘Expertise’ (Rijkzwaan, Merksem, Belgium) was sown either in Jiffy-7® (Jiffy Products International BV, Zwijndrecht, the Netherlands) in a 51 well tray or Growcoons® (150M-H39, Klassman-Deilman, Brugge, Belgium) in a 150 well tray (see Table 1). The nutrient solution consisted of 12.99 mmol NO_3_^-^, 1.35 mmol H_2_PO_4_^-^, 2.11 mmol SO_4_^2-^, 6.74 mmol K^+^, 4.13 mmol Ca^2+^, 1.69 mmol Mg^2+^ and 0.02 g L^-1^ micronutrients (Chelal Flor NF, BMS Micronutrients N.V., Bornem, Belgium).

**Table 1.**
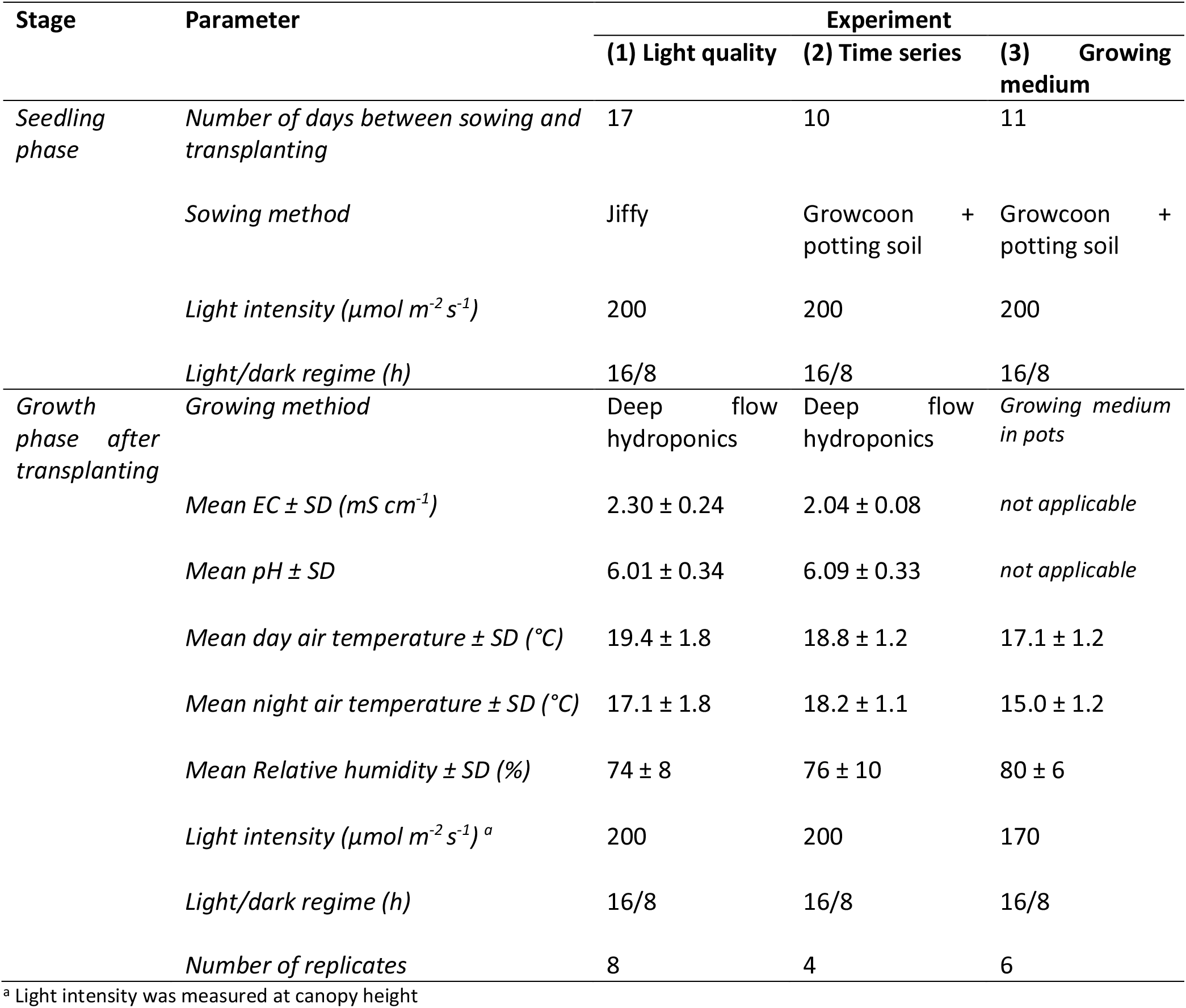
Experimental conditions.

### Light quality effect on root exudation rates

Rooted seedlings were transplanted 17 days after sowing to a deep flow hydroponic culture room with six plants per tray (30 x 40 x 12 cm, w x l x h) containing 3.5 L nutrient solution aerated with an aquatic stone (Supplemental figure 1). EC, pH, air temperature, and relative humidity were recorded (Table 1). From day 15 to day 19 after transplanting (DAT), plants were temporally moved to a growth room with cool-white fluorescent lamps (Osram T8 Lumilux 36W-840, Osram, Munich, Germany) 102 µmol m^-2^ s^-1^. Treatments consisted of three light qualities with eight trays (replicates, 24 trays in total): full-spectrum light from NS1/NS12 lamps (white; Valoya, Helsinki, Finland), 100% blue LED light at 450 nm (B200; Philips, Eindhoven, the Netherlands) and 100% red LED light at 660 nm (R200; Philips, Eindhoven, the Netherlands; Supplemental Figure 2). Samples were collected at 27 days after transplanting as described in ‘sampling’.

### Time effect on root exudation rates

Seedlings were transplanted in the deep flow hydroponic culture room under white, B200 and R200 lights (as specified in ‘light quality effect on root exudation rates’) with 4 replicates per treatment (Supplemental figure 3). EC, pH, air temperature and relative humidity were recorded (Table 1). Samples were collected 10, 17 and 24 days after transplanting as described in ‘sampling’.

### Growing medium effect on root exudation rates

The growing media consisted of potting soil (20% dry matter content, 10% organic matter, pH range from 4.5 to 7.5, EC of 0.75 mS cm^-1^, NPK 14-16-18 with spore elements 0.5 kg m^-3^, Aveve, Leuven, Belgium) mixed with perlite (1-3 mm, Agaris, Ghent, Belgium) at ratios specified in Table 2, resulting in mixtures with a different coarseness.

**Table 2.** Growing media treatments in the substrate experiment.

| Treatment | Potting soil (%) | Perlite (%) |
| --- | --- | --- |
| 100ps | 100% | 0 |
| 80ps | 80% | 20% |
| 50ps | 50% | 50% |
| 20ps | 20% | 80% |
| 0ps | 0% | 100% |

Ten-day old seedlings were transplanted to 13 cm pots containing the corresponding soil/perlite mixture and cultured in an ebb and flood system (Supplemental figure 4). Tables were flooded for 10-minutes with nutrient solution (as specified in ‘plant material and growth conditions’) every few days depending on soil moisture monitoring using a W.E.T. sensor (Delta-T Devices, Cambridge, UK). Pots were watered by flooding tables at moisture content < 30%. Air temperature and relative humidity were as shown in Table 1. Upon transplantation to the 13 cm pots, the three light regimes were applied: white, B170 and R170. The light spectra were used as specified in ‘light quality effect on root exudation rates’ (Supplemental Figure 2), but at 170 instead of 200 µmol m^-2^ s^-1^. Samples were collected 9 and 10, 16 and 17, and 23 and 24 days after transplanting as described in ‘sampling’.

### Sampling

In the light quality experiment, exudate was collected from a randomly selected plant per tray (eight trays (replicates)/treatment) at 27 days after transplanting. In the time series experiment (four trays (replicates)/treatment) root exudate was samples at 10, 17 and 24 days after transplanting. In the growing medium experiment (six pots (replicates)/treatment), root exudate was sampled at 9 and 10, 16 and 17 and 23 and 24 days after transplanting. Before collecting exudate, roots were washed under running tap water to remove the soil. Roots were submerged several times in distilled water and once in the sampling solution to reduce initial shock effects on the root exudation rate. Exudate was collected by submerging the roots in 100 ml MQ with 0.01 g L^-1^ Micropur for 2-3h. The plants with the sampling solution were placed under the different lights during sampling, which always took place between five to eight hours after the onset of light. During the sampling, the roots in the sampling solution were wrapped in aluminium foil. Micropur was added to inhibit microbial decomposition of exuded metabolites during the sampling period (Otxandorena-Ieregi *et al*., 2024; Santangeli *et al*., 2024). Plants were removed from the sampling solution after exudate sampling. Extracted exudate was filtered using a 0.2 µm cellulose acetate filter to remove microbial contaminants, then freeze-dried and stored at -20°C. After exudate sampling, the root tissue was dried at 70°C until even weight to record dry weight (DW).

### Exudate analysis

Total exuded C as µmol C per g root DW per hour was determined by measuring the absorbance of 250 μl root exudate sample at 260 nm with potassium phthalate for the calibration following Oburger et al. (2022) with a Tecan plate infinity reader (Tecan Group Ltd., Männedorf, Switzerland).

Total exuded carbohydrates were determined with the anthrone reaction as described by Hansen and Møller (1975). First, 100 μl freeze-dried root exudate sample that was resuspended in 4 ml MQ water was mixed with 200 72% H_2_SO_4_. This was mixed with 400 μl anthrone reagent (2g antrone/L 72% H_2_SO_4_) and heated for 15 minutes at 95°C. After cooling to room temperature, absorbance was determined at 625 nm with a Tecan plate infinity reader (Tecan Group Ltd., Männedorf, Switzerland). Glucose was used for the calibration curve. Total carbohydrate exudation rate was reported as µmol glucose equivalent (gluc. equiv.) per g root DW per hour.

Total phenolics were analyzed using the Folin-Ciocalteu method (Liu *et al*., 2007). First, 50 μl freeze-dried root exudate sample that was resuspended in 1 ml MQ water was added to 475 μl 0.25M Folin&Ciocalteu reagent. Samples were incubated in the dark for two minutes before 475 μl 1M Na_2_CO_3_ was added. Samples were incubated for 60 minutes in the dark before measuring absorbance at 765 nm with a Tecan plate infinity reader (Tecan Group Ltd., Männedorf, Switzerland). Total phenolic exudation rate was expressed as µmol gallic acid equivalents (GAE) per g root DW per hour.

### Statistical analysis

Statistical analyses were done using R (R Core Team, 2023). The data were tested for normality with the Shapiro-Wilk test and homogeneity of variances with Levene’s test (Fox and Weisberg, 2019). Normally distributed data from the light quality experiment was tested for differences between treatments with a one-way analysis of variance (ANOVA) with light as main factor. Normally distributed data from the time series experiment and the growing medium experiment was tested with a multifactor ANOVA with light and time, or light, time, and medium as main factors, respectively. Tukey’s honest significance difference test was used as a post hoc test (de Mendiburu, 2021). If data were not normally distributed (indicated under each figure) or had no homogeneity of variances, the non-parametric alternative Kruskal-Wallis test was used for the one-way ANOVA and a Dunn’s test with multiple comparisons with Benjamini-Hochberg p-adjustment (Kassambara, 2021). As a non-parametric alternative for the two-factor ANOVA a Scheirer Ray Hare test (Mangiafico, 2023) was used with a Dunn’s test with multiple comparisons with Benjamini-Hochberg p-adjustment (Kassambara, 2021). As a non-parametric alternative for the three-factor ANOVA an ANOVA with permutations (Wheeler and Torchiano, 2016) was used. Two exudate samples from the light quality experiment of plants grown under white light contained a noticeable amount of soil particles, which resulted in outliers, and were consequently removed from the dataset.

## RESULTS

To separate the effect of light quality on photosynthesis and root exudation, pure blue and pure red LED light intensities were adjusted to reach identical photosynthetic active radiation levels. Deep flow hydroponic lettuce was irradiated with white, red, and blue LED light. Total C exudation was determined as a measure of net exudation rate. The total C exudation was similar for plants grown in red, blue and white light, indicating that in this culturing system root exudation is not controlled by light quality (Figure 1A).

**Figure 1.**
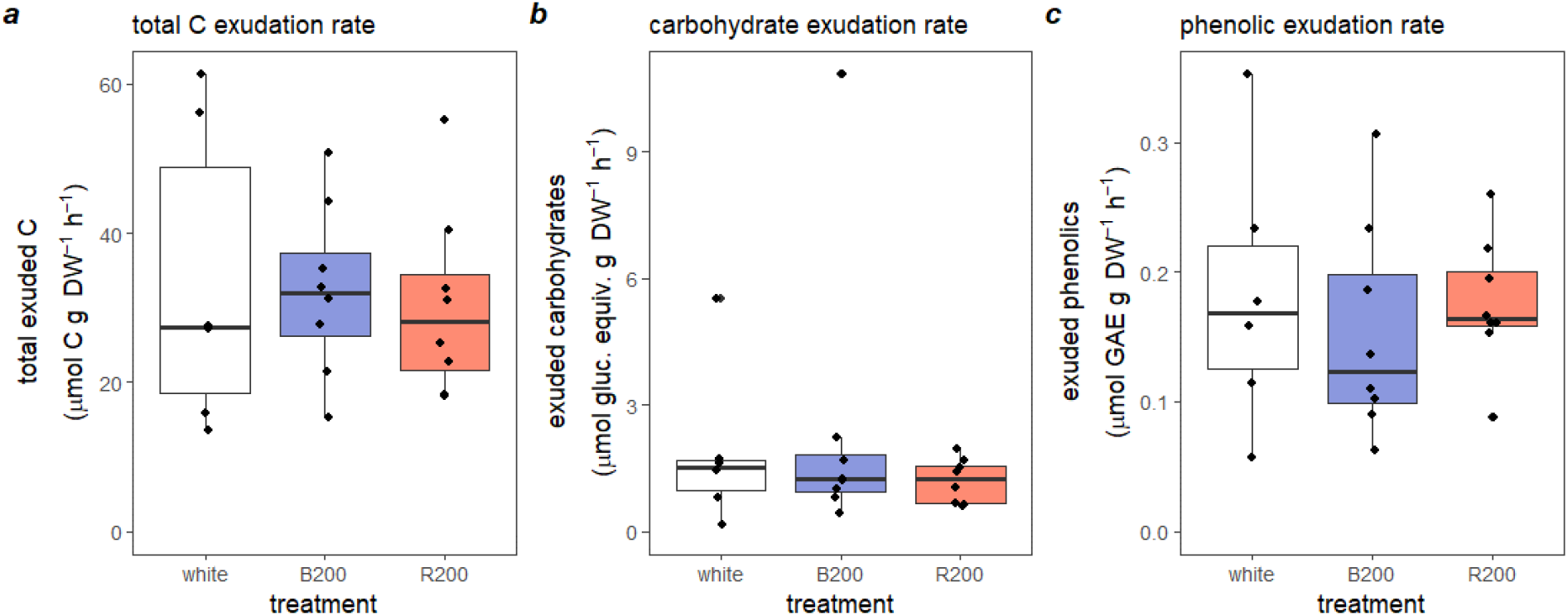
Boxplot of A) total carbon exudation rate (µmol C g root DW^-1^ h^-1^), B) total carbohydrate exudation rate (µmol glucose equivalents g root DW^-1^ h^-1^) and C) total phenolic exudation rate (µmol gallic acid equivalents (GAE) g root DW^-1^ h^-1^) of lettuce plants grown in DFT in Jiffy’s under white light, 100% blue light (b200) and 100% red light (r200), measured after 27 days of treatment. Significant differences were tested with a Kruskal-Wallis test and are indicated with letters. Absence of letters indicates no significant differences (n=8).

The analysis was further refined by measuring the total carbohydrate and the total phenolics in the root exudate (Figure 1B, C) as carbohydrates and phenolic compounds are considered major determinants of the plant-associated microbiome (Zwetsloot *et al*., 2018; Mavrodi *et al*., 2021). Under white, blue, and red light, we observed similar levels of carbohydrates and phenolics, indicating that light quality did not lead to strong alterations in root exudate composition (Figure 1B, C).

In view of the absence of a significant impact of light quality on exudation, we suspected that the plants were late in the vegetative growing cycle and that this would have led to very low exudation rates (Hamlen *et al*., 1972; Groleau-Renaud *et al*., 1998). Using the same light conditions, we therefore sampled at earlier time points and determined the total C exudation rates. Exudation rates significantly (p < 0.05; Table 3) decreased by 78% between 10 and 24 days after exposing the plants to the different light qualities (Figure 2). Remarkably, we observed no difference in exudation rate under blue, red, or white light growing conditions (Figure 2; Table 3).

**Table 3.** *P*-values of the Scheirer Ray Hare test on total exuded C in the time series experiment. * indicate significant *p*-values (*p* < 0.05).

| Variable | $p$-value |
| --- | --- |
| Light | 0.75236 |
| DAT | 0.00005 * |
| Light * DAT | 0.98000 |

**Figure 2.**
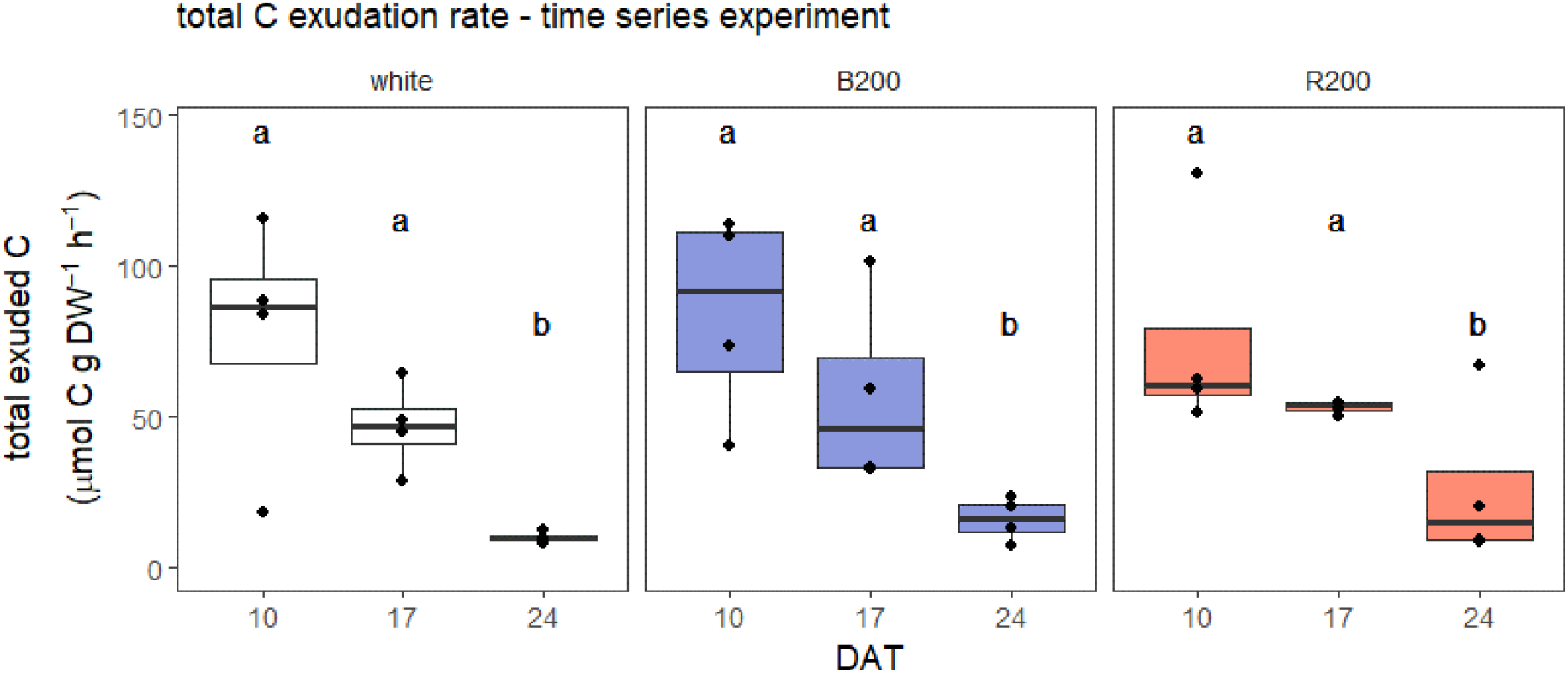
Boxplot of total carbon exudation rate (µmol C g root DW^-1^ h^-1^) of lettuce plants grown in deep flow hydroponics under white light, 100% blue light (B200) and 100% red light (R200) measured at 10, 17 and 24 days after transplanting (DAT). As data was not normally distributed, significant differences were tested with a Scheirer Ray Hare test as shown in Table 3 (n=4).

As growing medium structure has been shown to influence root exudation (Sasse *et al*., 2020), we explored the possibility that root architecture interacts with light quality on root exudation. To evaluate the effect of root architecture, lettuce was grown in different ratios of potting soil and mineral growing media with bigger particles. The total C exudation rates were measured from plants grown in these substrate mixtures (Figure 3). Exudation decreased over time (Figure 3). The highest decrease was between 9/10 and 16/17 days after transplanting (39% for the plants that decreased). The change in root exudation rates was smaller between 16/17 and 23/24 days after transplanting (2% increase and 35% decrease on average across all growing media). The highest variation was found at the first sampling moment (range, i.e. lowest value subtracted from the highest value, is 1456 µmol C g DW^-1^ h^-1^) and the lowest variation at 23/24 days after transplanting (range is 544 µmol C g DW^-1^ h^-1^). In contrast with the previous experiment, the total C root exudation rates depended on the light quality. Statistical analyses revealed a significant (p < 0.05) interaction between light quality, growing medium and chronological age (Table 4). For plants grown in 100% potting soil, the means of the different light conditions deviated less from each other than the means in the other growing media conditions. At the first sampling moment (9/10 DAT), white or blue light resulted in the highest total C exudation rate, with exception of plants grown in 50% potting soil. In this 50% growing medium, red light had a higher median than blue and white light. At the second sampling moment (16/17 DAT), light quality only had an effect at 80% potting soil. Red light causes a lower C exudation rate. At the last sampling moment (23/24 DAT), red light was associated with the lowest C exudation rate albeit that the difference was substantial when using 100% potting soil. Moreover, total C exudation rate shows an optimum at the last time point for the percentage in potting soil, with minimum values at 50% and 20% potting soil.

**Table 4.** *P*-values of the ANOVA test with permutations on total exuded C in the growing medium experiment. * indicate significant *p*-values (*p* < 0.05).

| Variable | *p*-value |
| --- | --- |
| Light | 0.0425 * |
| Growing medium | 0.6879 |
| Light: growing medium | 0.2352 |
| DAT | $<2\text{e}^{-16}$ * |
| Light: DAT | 0.6062 |
| Growing medium: DAT | 0.3482 |
| Light: growing medium: DAT | 0.0164 * |

**Figure 3.**
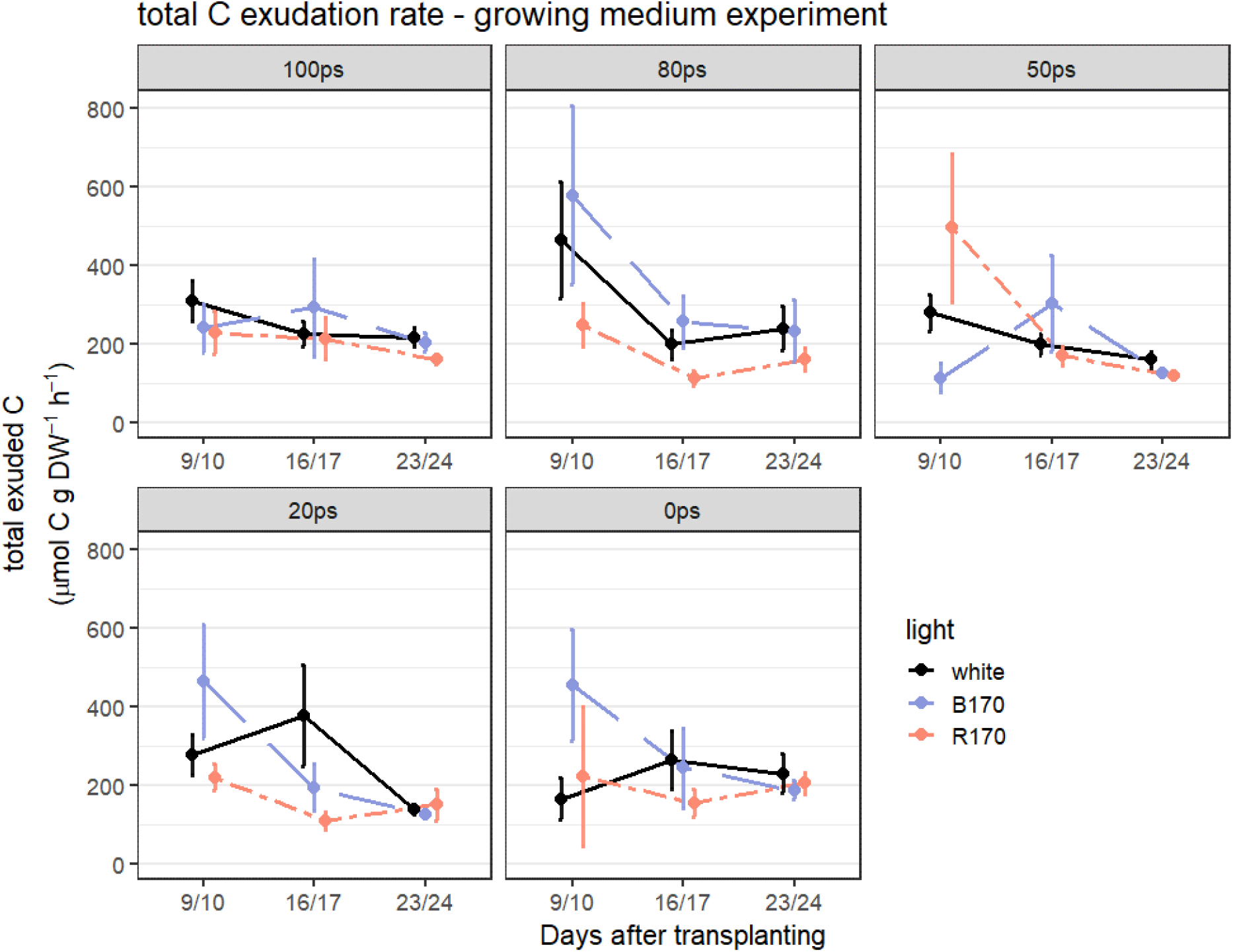
Mean total carbon exudation rate ± SE (µmol C g root DW^-1^ h^-1^) of lettuce plants grown under 170 μmol m^-2^ s^-1^ white light, 100% blue light (B170) and 100% red light (R170) measured at 9/10, 16/17 and 23/24 days after transplanting (DAT) in different growing media ranging from 100% potting soil (100ps) to 0% potting soil (0ps), combined with perlite (n=6).

Next, we asked how light quality influences the composition of root exudates. For this, the contribution of carbohydrates in the observed variation in root exudation was analyzed with respect to growing medium and culturing time. Interestingly, the interaction effect of light quality and sampling time was significant (p < 0.05; Table 5). Plants grown under white light showed an increase in carbohydrate exudation rates over time (Figure 4). The mean increased from 4.78 to 6.70 to 7.73 µmol glucose equivalents g DW^-1^ h^-1^. Blue (B170) and red (R170) light treatments resulted in the lowest carbohydrate exudation rates at 16/17 days after transplanting compared to the other sampling moments. Red light resulted in a higher biomass than plants grown under white or blue LED (Supplemental Figure 5). The application of red light also increased the carbohydrate content which in the growing media experiment correlated with a decreased exudation rate (p = 0.047; Supplemental figure 6).

**Table 5.** *P*-values of the ANOVA test with permutation on total carbohydrate exudation rate in the growing medium experiment. * indicate significant *p*-values (*p* < 0.05).

| Variable | $p$-value |
| --- | --- |
| Light | 0.1765 |
| Growing medium | 0.0700 |
| Light: growing medium | 0.1423 |
| DAT | $< 2e^{-16}$ * |
| Light: DAT | $< 2e^{-16}$ * |
| Growing medium: DAT | 0.4849 |
| Light: growing medium: DAT | 0.3464 |

**Figure 4.**
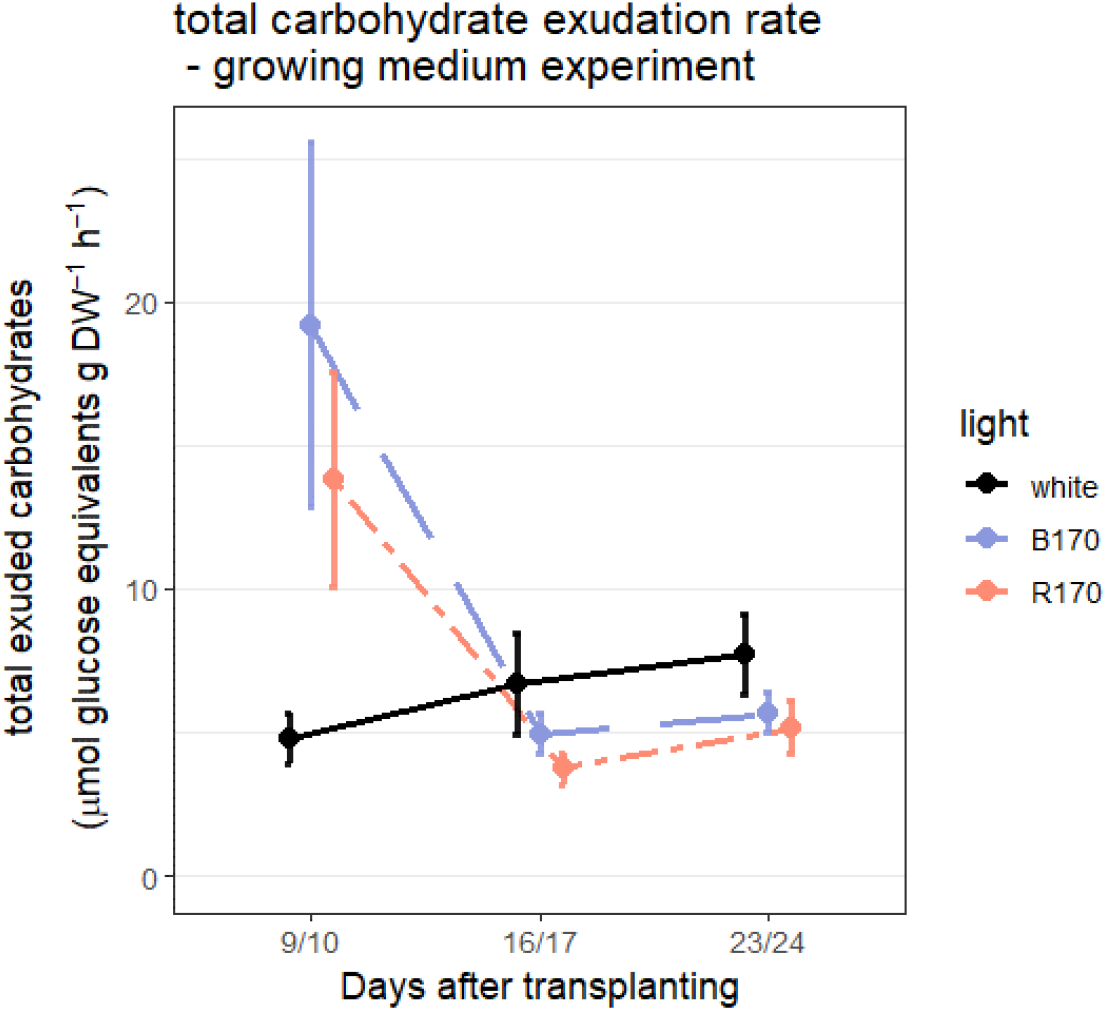
Mean carbohydrate exudation ± SE rate (µmol glucose equivalents g root DW^-1^ h^-1^) of lettuce plants grown dependent on light conditions (170 μmol m^-2^ s^-1^ white: full spectrum light, B170: monochrome blue light at 450 nm, R170: monochrome red light at 660 nm). Data are pooled for substrate after significant differences were tested with an ANOVA test with permutations as shown in Table 5, as data was not normally distributed (n=6).

Besides total carbohydrate exudation rates, total phenolic exudation rates were also determined. Total phenolic exudation rates were significantly (p < 0.05) influenced by the interaction between growing medium and culturing time (Table 6). At the first sampling moment, the variation was much higher (6.11 µmol GAE g DW^-1^ h^-1^) than at the last sampling moment (2.36 µmol GAE g DW^-1^ h^-1^) (Figure 5). At the first sampling moment, the percentage of potting soil had the strongest impact on the total phenolic exudation rates. The most extreme growing media resulted in the highest mean for exudation rates (2.59 and 2.64 µmol GAE g DW^-1^ h^-1^), while the lowest mean is found at 50% potting soil (1.19 µmol GAE g DW^-1^ h^-1^).

**Table 6.** *P*-values of the ANOVA test with permutation on total phenolic exudation rate in the growing medium experiment. * indicate significant *p*-values (*p* < 0.05).

| Variable | *p*-value |
| --- | --- |
| Light | 0.1452 |
| Growing medium | < 2e<sup>-16</sup> * |
| Light: growing medium | 0.3338 |
| DAT | < 2e<sup>-16</sup> * |
| Light: DAT | 0.1018 |
| Growing medium: DAT | < 2e<sup>-16</sup> * |
| Light: growing medium: DAT | 0.2940 |

**Figure 5.**
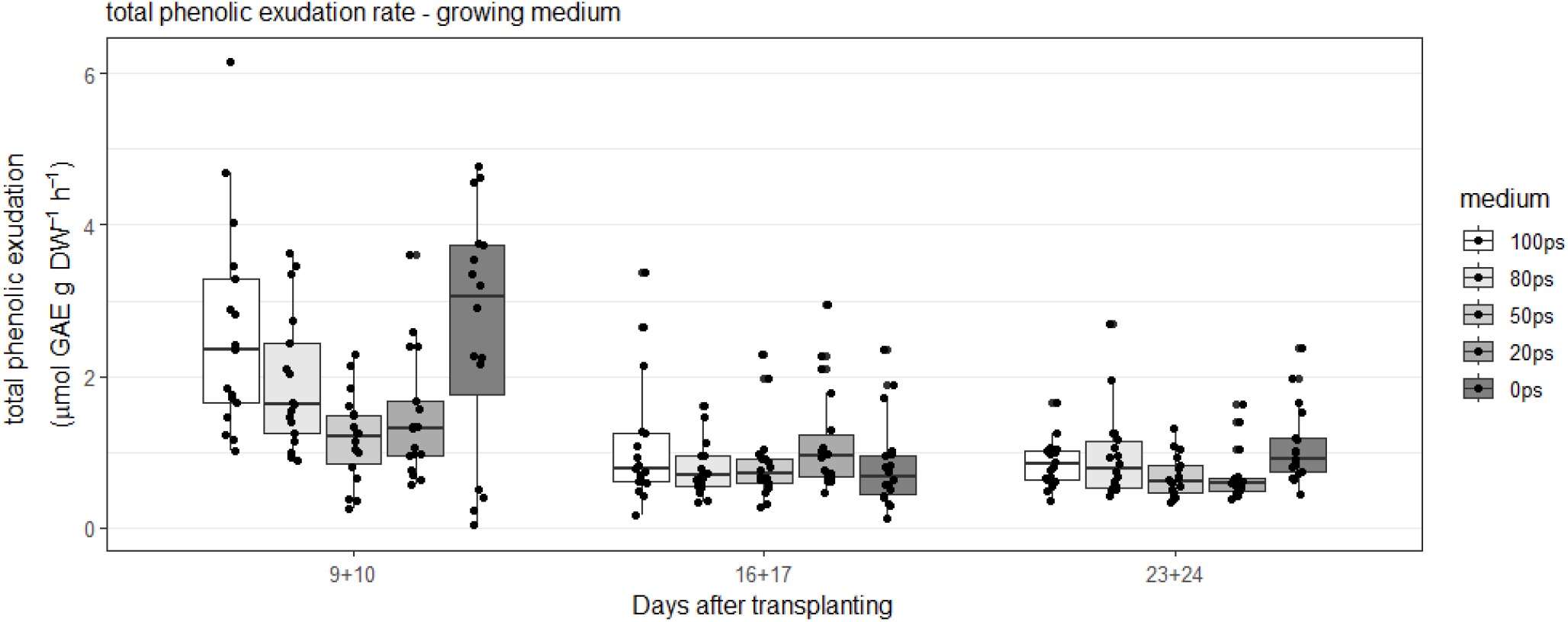
Boxplots of total phenolic exudation rate (µmol gallic acid equivalents (GAE) g root DW^-1^ h^-1^) of lettuce plants measured at 9/10, 16/17 or 23/24 days after transplanting grown in a gradient from 0% to 100% potting soil (ps). Data are pooled for light treatment, after significant differences were tested with an ANOVA test with permutations as shown in Table 6, as data was not normally distributed (n=6).

## DISCUSSION

Root exudation is often regarded of lesser importance in hydroponic cultures compared to soil-grown plants because rough estimations indicate lower exudation rates (Oburger *et al*., 2013). However, microbiome analyses of the rhizosphere of hydroponically grown roots show strong variations which may affect yield and tolerance to pathogens (Van Gerrewey *et al*., 2024). In this work, we asked whether root exudation can be controlled by light conditions. To achieve this, we set out a series of experiments with soilless grown lettuce and analyzed the root exudate derived from plants grown under white, blue, and red light.

The total C released by the root system in hydroponics is highly dynamic corroborated by the observation that lettuce root exudation strongly reduces as the plant ages (Hamlen *et al*., 1972; Groleau-Renaud *et al*., 1998). In the time series experiment, total C exudation rate declined 78% between 10 and 24 DAT and in the growing media experiment it dropped 44% between 9/10 and 16/17 DAT. These results are in line with other studies that show a decline in root exudation with plant age in several crops, including *Sorghum bicolor, Crotalaria juncea, Elucina coracana, Solanum lycopersicum* (Balasubramanian and Rangaswami, 1969), *Oryza sativa* (Zang *et al*., 2019), *Zea mays* (Groleau-Renaud *et al*., 1998; Santangeli *et al*., 2024) and *Medicago sativa* (Hamlen *et al*., 1972). The physiological basis for a decrease in total C exudation rates is hypothesized to be linked to an increase in suberization of the physiologically older root sections (Ranathunge and Schreiber, 2011) that physically prevents the secretion of small molecules.

Culturing time not only affected the total C exudation rate, but also the total phenolics exudation rate. The total phenolic exudation rate was highest in the 100% potting soil and perlite in the first culturing phase (9/10 DAT). Growing media, or soils have been shown to influence exudation through particle size or mechanical impedance (Barber and Gunn, 1974; Groleau-Renaud *et al*., 1998). This can alter root morphology, though not one morphological parameter seems to thoroughly explain the observed changes (Groleau-Renaud *et al*., 1998). Another way through which soil can change root exudation, is through metabolite adsorption (Sasse *et al*., 2020). Clay absorbs a variety of metabolites, including charged compounds, amino acids and sugars, changing the exudation profile. In addition to structural differences between these materials, perlite is expanded volcanic glass for which we assume that the microbial load is much lower than for the potting soil (Bardopoulou *et al*., 2001; Alsanius and Wohanka, 2019). It is hence conceivable that the impact of light on exudation depends on the presence of microbes in the growing medium. Korenblum et al. (2020) showed that a decrease in microbial species, led to a change in the root exudate profile of *Solanum lycopersicum*. Mainly secondary metabolites changed in abundance, such as acyl sugars, alkaloids and hydroxycinnamic acid conjugates. Moreover, lettuce was found to alter the exudation rates of benzoic acid and lauric acid in the presence of specific pathogens (Windisch *et al*., 2017). Moreover, root exudation may also be changed by nutrient availability (Sardans *et al*., 2023), though in the current experiment its impact cannot be proven. The impact of growing media could therefore be direct through particle size or metabolite absorption, or indirect through the root-associated microbiome.

Though our first set of experiments indicated that light quality did not affect root exudation rates, surprisingly, light quality changed carbohydrate exudation rates when lettuce was grown in growing media. The impact of light quality on total C exudation was most pronounced at the early growth phase and converged towards the end of the lettuce culture. Culturing lettuce under white LED showed a relatively carbohydrate low exudation rate that steadily increased over time, whereas monochrome red or blue showed an initially high level of exudation that rapidly declined after the first sampling time point. As red light resulted in a higher biomass than plants grown under white or blue LED, the difference in exudation is unlikely linked with the efficiency of photosynthesis. The application of red light also increased the carbohydrate content which in the growing media experiment correlated with a decreased exudation rate (−0.21, p = 0.047). A previous study has shown that a high carbohydrate leaf content correlated with a low carbohydrate exudation rate (Barber and Gunn, 1974). The impact of red and blue light on exudation is therefore not likely through the accumulation of carbohydrates, but mediated through other light controlled processes.

In 100% potting soil, light quality did not differentially affect total C exudation, but for the other substrate compositions it did, especially during the first phase of the culture (time point 9/10 DAT). Blue light conditions were associated with a high total C exudation, except for the substrate mix with 50% potting soil where red light stimulated exudation. In general, the data indicate that light quality can influence exudation at an early developmental stage and as the lettuce plant ages, light no longer is a determining factor.

The effect of light quality on carbohydrate exudation was the strongest in pure potting soil as opposed to pure perlite (Supplemental Figure 2). Growing media rich in perlite, however, showed a tendency to secrete more carbohydrates independent from the light used. These surprising results indicate that growing media composition is an important factor that determines exudation rate adaptation in function of the light conditions.

We speculate that the presence of a chemical or the microbial context is important for the plant to create exudate homeostasis. Growing media (Van Gerrewey *et al*., 2024) and soils (Schreiter *et al*., 2014) have been shown to strongly influence the root-associated microbiome. Therefore, the relation between culturing method and light quality on exudation may be mediated by the microbiome. Future studies should explore this possibility. That light quality plays a role in establishing an appropriate exudation rate and may therefore be related to the establishment of the root microbiome. Under low microbial load, the root system may be subjected to fewer feedback loop regulation which allows the secretion machinery to be more sensitive to other external factors such as light quality.

## CONCLUSIONS

Plant growth promoting bacteria (PGPB) improve the microbiome of a plant, and therefore indirectly the plant itself. However, the effects of PGPB can be variable (Consentino *et al*., 2022). A more sustainable way to improve the plants microbiome, would be if plants themselves could steer it. Root exudates play a role in this (Mondal *et al*., 2023), and growers can ideally influence these exudates through environmental factors. In this study, light conditions, substrate and chronological age are shown to be important determinants of root exudation rates. However, light quality interacts with culturing medium on root exudation of carbohydrates. Phenolics, on the other hand, were mainly influenced by the growing medium itself. Thus, light quality changes root exudation of soilless grown lettuce depending on culturing method and plant chronological age.

## ACKNOWLEDGEMENTS

The authors wish to thank Chiara Degli Eposti, Stefano Triolone, Liv Van Haver, Xinquan Hu and Tanguy Van de Wege for their help with sampling root exudates. We thank Christophe Petit for his technical support. We are grateful for the support during the growing medium experiment from Chiara Degli Eposti, and for the help with the laboratory analysis from Jana Decuyper, Liv Van Haver and Chiara Degli Eposti. We wish to thank Charlotte Grootaert for freeze drying samples. This research has benefitted from a statistical consult with Ghent University FIRE (Fostering Innovative Research based on Evidence).

## AUTHOR CONTRIBUTIONS

DG, ED, MCVL, BDH: conceptualization; BDH: investigation; EO: methodology root exudate analysis; BDH: data curation; BDH: formal analysis; DG, ED, MCVL: resources; DG, ED, MVCL: supervision; BDH: visualization; BDH: writing – original draft preparation; BDH, EO, MCVL, ED, DG: writing – review & editing.

## DATA AVAILABILITY

Statement: All primary data to support the findings of this study are openly available in Zenodo at [https://doi.org/10.5281/zenodo.11386429] .

## ABBREVIATIONS

GAE: gallic acid equivalents
GC-MS: gas chromatography-mass spectroscopy
EC: electrical conductivity
DAT: days after transplanting
B200: 100% blue light (peak at 450 nm) at 200 µmol m^-2^ s^-1^
R200: 100% red light (peak at 660 nm) at 200 µmol m^-2^ s^-1^
B170: 100% blue light (peak at 450 nm) at 170 µmol m^-2^ s^-1^
R170: 100% red light (peak at 660 nm) at 170 µmol m^-2^ s^-1^
ps: potting soil
DW: dry weight
PGPB: plant growth promoting bacteria

