## Supplemental Data for "Light and substrate composition control root exudation rates at the initial stages of soilless lettuce cultivation"

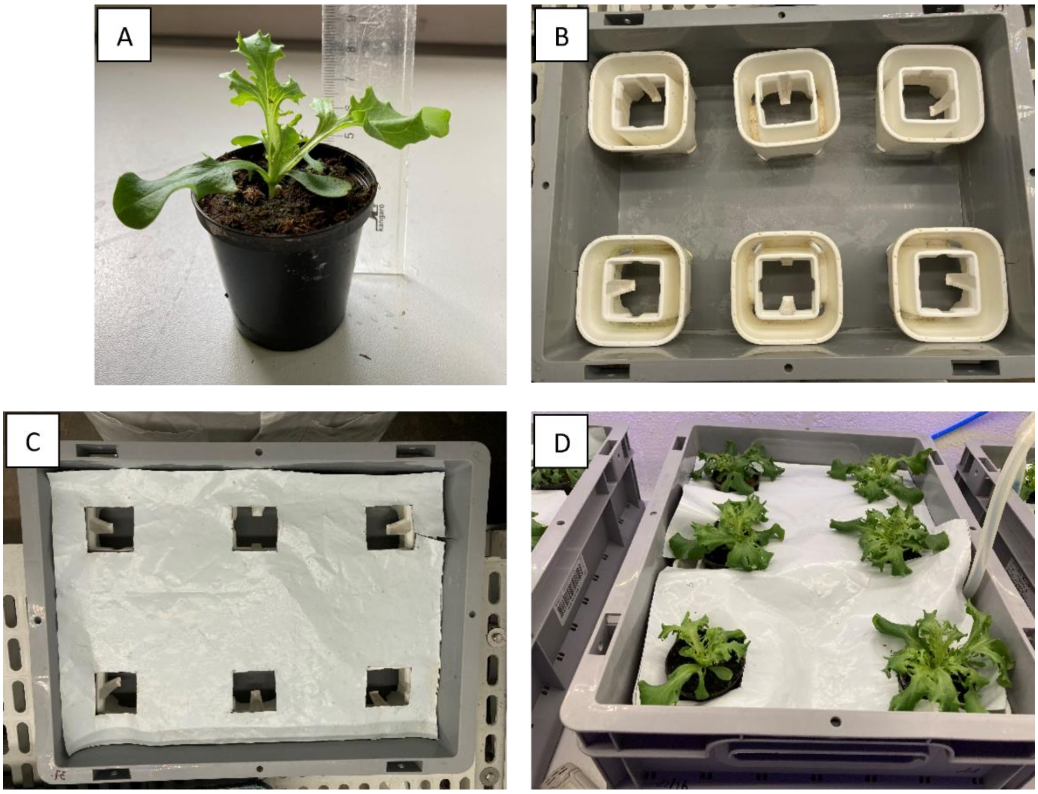


**Supplemental figure 1.** Details of the set up of the light quality experiment. A) a young plant grown in Jiffy, with the outer net removed. A plastic cover is used to hold the soil together, with a hole in the bottom to allow the roots to grow. B) The container (30 x 40 x 12 cm, w x b x h) with the plant holders to keep the roots partially in the nutrient solution. C) As B, but with a plastic foil to minimize light reaching the roots. D) The plants during the experiment. The tube on the right side of the picture is connected to an aeration stone inside the water to have sufficient oxygen levels in the nutrient solution.

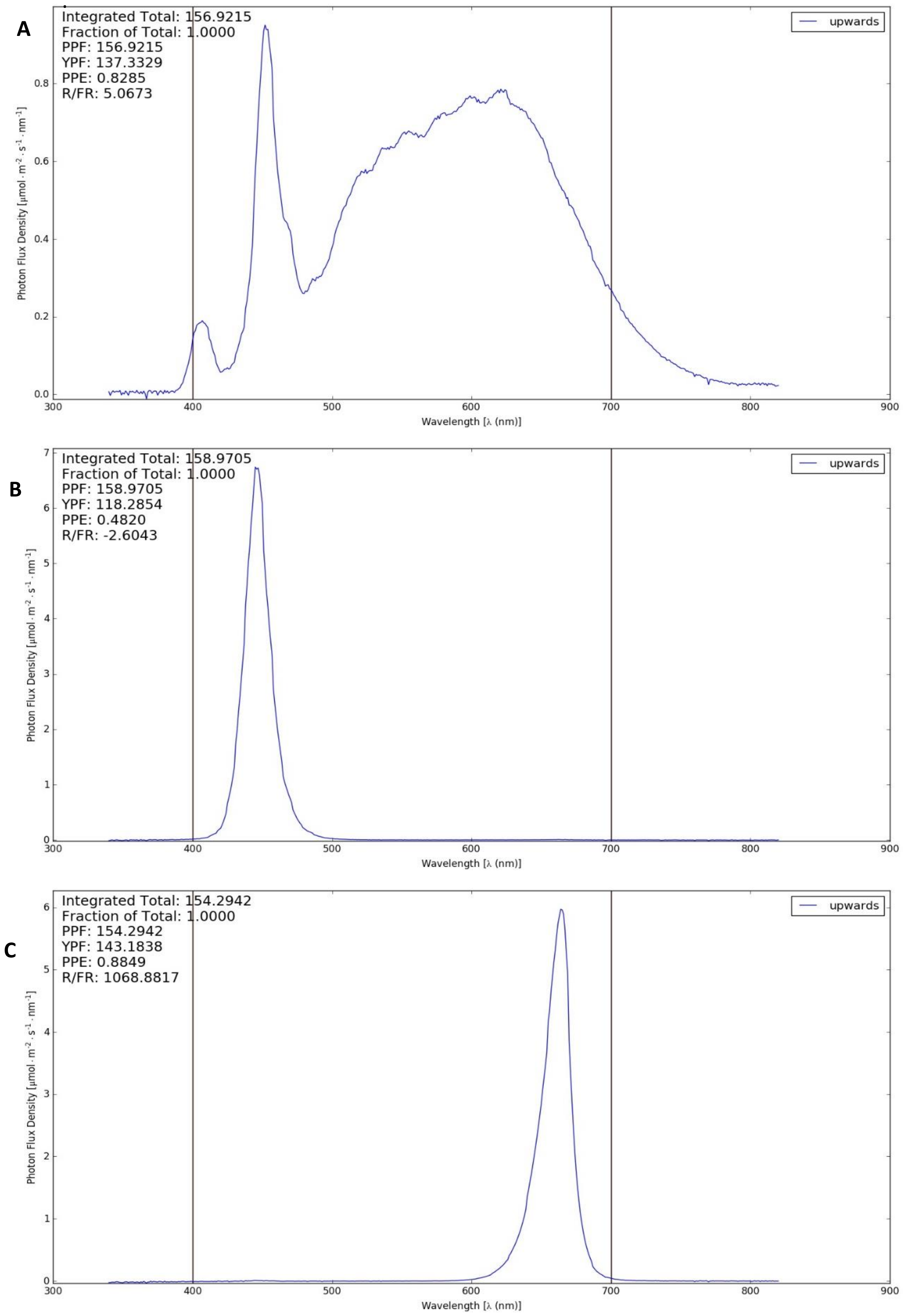


**Supplemental Figure 2.** Light spectra of the three light conditions. A) white: full spectrum light, B) blue: peak at 450 nm, C) red: peak at 660 nm.

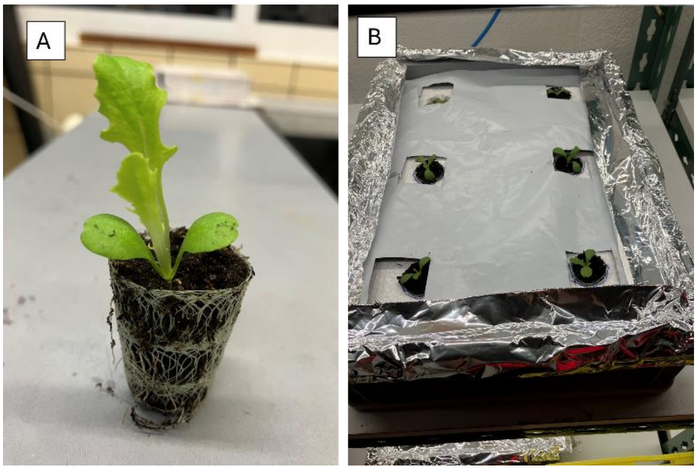


**Supplemental figure 3.** Details of the setup of the time series experiment. A) A young plant in a grow coon with potting soil at 10 days after transplanting. B) Grow coons with plants in styrofoam with a horticultural foil to cover the roots from the light. The aluminium foil was a temporal solution and replaced with strips of horticultural foil.

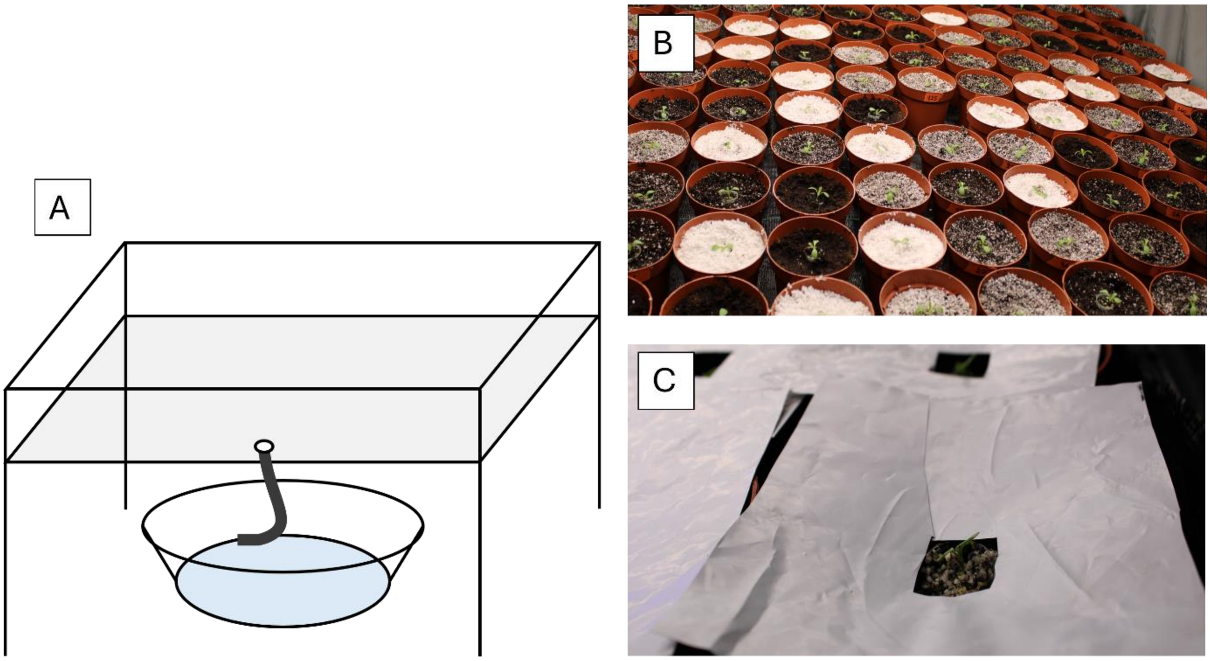


**Supplemental figure 4.** Details of the setup of the growing media experiment. A) A schematic drawing of a ebb and flood table. Underneath the table a barrel was placed and filled with nutrient solution. A pump and pipe allowed to flood the table at the desired moments. Water could stay on the table as long as needed by closing of the drain. When plants had been flooded for long enough (usually 10 minutes), the table was drained through the same pipe. B) Image of the plants just after planting the grow coon plugs in the pots. C) The soil was covered with horticultural foil to prevent reflection effects of the different coloured growing media.

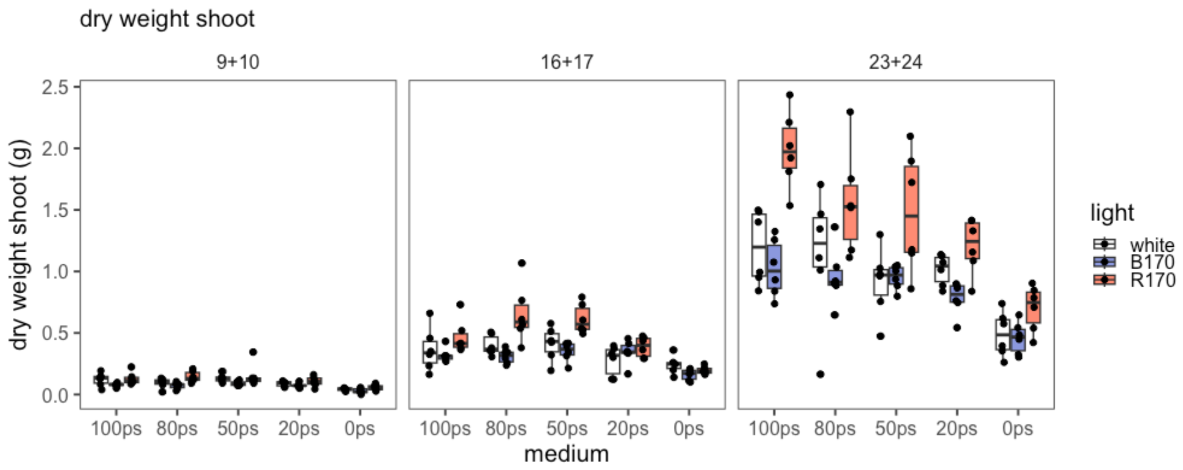


**Supplemental Figure 5.** Boxplot of the shoot dry weight (g) of lettuce plants grown under white light, 100% blue light (B170) and 100% red light (R170) measured at 9/10, 16/17 and 23/24 days after transplanting (DAT) in different growing media ranging from 100% potting soil (100ps) to 0% potting soil (0ps), combined with perlite.

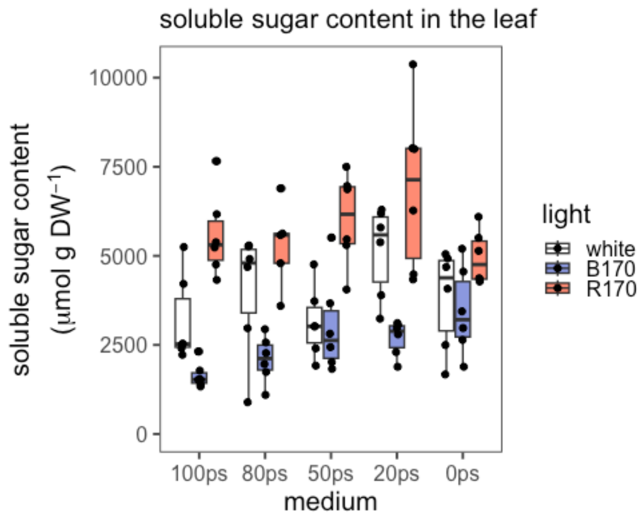


**Supplemental Figure 6.** Boxplot of the soluble sugar content (glucose, fructose and sucrose; µmol g $DW^{-1}$) of lettuce plants grown under white light, 100% blue light (B170) and 100% red light (R170) measured at 9/10, 16/17 and 23/24 days after transplanting (DAT) in different growing media ranging from 100% potting soil (100ps) to 0% potting soil (0ps), combined with perlite.
